# Enhanced protein precipitation with ammonia enables rapid, universal extraction of oligonucleotides for bioanalysis

**DOI:** 10.1101/2025.04.24.650521

**Authors:** Afrand Kamali, Juili D. Shelke, Ethan J. Sanford, Michael M. Hayashi, Natthamon Chaisakhon, Robert V. Kolakowski, Sanyogitta Puri, Guangnong Zhang

**Affiliations:** Dicerna Pharmaceuticals, a Novo Nordisk Company, 65 Hayden Ave, Lexington, MA 02421, United States; Nucleic Acid Delivery, Formulations & Bioanalysis. Novo Nordisk, Lexington Massachusetts 02421, United States; Nucleic Acid Chemistry. Novo Nordisk, Lexington Massachusetts 02421, United States

## Abstract

Extracting oligonucleotides from biological matrices for mass spectrometry (MS) using current methodologies is time-consuming and costly. Accelerating the discovery, characterization, and development of oligonucleotides (ONTs) requires a universal method that is low-cost, efficient, without requiring extensive method development for each new test article. Protein precipitation using organic solvents meets these criteria for small molecules but has historically failed for ONTs due to poor recovery arising from their tendency to co-precipitate with proteins. This study introduces an effective approach utilizing small amines dissolved in organic solvents to significantly boost extraction recovery of ONTs from biological matrices .We first present the method’s development and then analytically qualify it for the bioanalysis of antisense oligonucleotides (ASOs) and small interfering RNAs (siRNAs) extracted from various tissues, using ion-pairing reverse phase (IPRP) liquid chromatography coupled to tandem MS and high resolution MS (HRMS) detection. This method, Enhanced Protein Precipitation (EPP), achieves near-complete recovery across multiple ONT classes from biological matrices without need of sample digestion, costly solid phase extraction plates, or custom-designed hybridization probes. The method has proven to be more versatile and sustainable than conventional approaches.

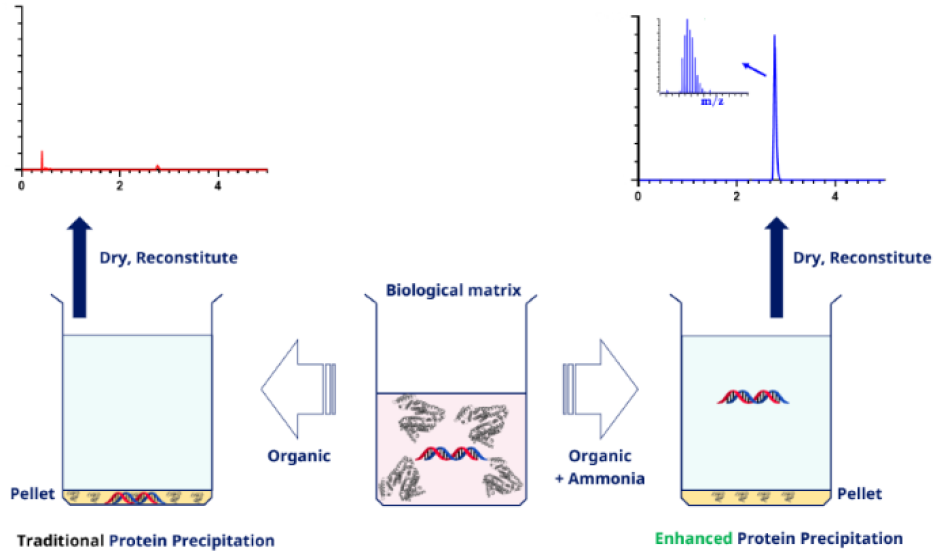

---

Oligonucleotide therapeutics are a swiftly advancing drug modality.^1–3^ ONTs include antisense oligonucleotides (ASOs), small interfering RNAs (siRNAs), and short hairpin RNAs (shRNAs) .^4, 5^ Successful discovery and development of any drug is contingent, in part, on the ability to measure, from biological samples, its absorption, metabolism, distribution. excretion (ADME) properties by bioanalytical assays.^6^ For ONTs, the predominant bioanalytical techniques used in the field are quantitative hybridization enzyme linked immunosorbent assay (hELISA),^7, 8^ quantitative polymerase chain reaction (qPCR) ^9^ or chromatographic separation coupled to either fluorescence detection or mass spectrometry.^10–15^Preference for MS-based analysis is attributed not only to its capability for quantification but also to its utility in metabolite identification, enabling differentiation between the absolute or relative concentrations of full-length products (FLPs) and their metabolites.^13^ Matrix proteins interact with nucleotides through a variety of interaction types involving diverse amino acid residues.^16-21^ The MS-based quantification of ONTs derived from biological matrices necessitates the elimination of interfering compo nents, particularly proteins, present within these matrices.^22, 23^ This challenge is also prevalent in the analysis of small molecule therapeutics; however, for this modality typically the introduction of organic solvents for precipitating proteins, commonly referred to as “protein precipitation” or “crashing”, serves as a straightforward extraction method.^24-26^ Since interaction be tween anionic backbone of ONTs and positively charged residues of proteins is not eliminated by organic solvents, addition of these solvents causes co-precipitation of ONTs with proteins, resulting in diminished recovery rates.^27, 28^ Contemporary extraction methods for ONTs employ strategies to mitigate interference from matrix proteins by using proteolytic enzymes for digestion often accompanied by use of lysis reagents; conducted at 55 °C or higher for durations ranging from 1 to 2 hours.^29,30^. Notable examples include solid-phase extraction (SPE), liquid-liquid extraction (LLE), and probe-based hybridization techniques prior to LCMS analysis. ^29-31^ While each of these techniques presents distinct advantages, they often entail substantial costs, labor-intensive procedures, and class-specific method development. Furthermore, the current strategies do not guarantee sufficient recovery for the constantly emerging novel conjugated ONTs.^29, 32, 33^ A simple extraction approach like those available for small molecules, protein precipitation, would positively impact the process of ONT drug discovery and development. Reports on using protein precipitation for ONTs are scattered and their limited success was primarily observed with specific classes of oligonucleotide chemistries.^29, 34, 35^ Interactions of amines with nucleotides and their polymers have a long history. Examples of leveraging these interactions for scientific purposes are use of amines for ion-pairing reversed phase (IPRP) chromatography of ONTs,^36-38^ stabilization of mRNA in lipid nanoparticles via ionizable lipids bearing amine groups,^39, 40^ presence of amines in sorbents of SPE.^41,42^

Further, covalent incorporation of amines attached to backbone of ONTs to neutralize negatively charged backbone for higher stability and better delivery of these therapeutics has been explored.^43^ Different types of amines were used by Khym^44^ serving as phase transfer catalyst in LLE for extraction of nucleic acids. Murray and Thompson used cetyltrimethylammonium bromide (CTAB) for extraction of high molecular weight DNA.^45^ The method and its modified versions^46^ utilizevariety of reagents that are not ideal for LCMS applications, like phenol and cesium chloride,-ethidium bromide, and are all lengthy procedures. Additionally, removal of quaternary amines like CTAB prior to LCMS analysis due to their positive charge and excess amounts is required, which is hard to achieve, adding more steps to the procedure. A few contemporary methods for bioanalysis of ONTs have reported benefits of using ammonium hydroxide in LLE or SPE.^23,47,48^ It is proposed that ammonium hydroxide disrupts the protein-nucleotide interaction and in case of lipophilic ONTs such as locked nucleic acids (LNAs) increases their solubility in aqueous phase.^47, 48^ The limitations imposed by LLE reagents like phenol, and interference from other hydrophilic extracts necessitates development of a method where interaction between amines and ONTs could result in a cleaner sample with a simpler workflow. Here we hypothesize that the recovery of ONTs with protein precipitation method can be enhanced by incorporating amines in organic solvents. In one report, by Johnson et al. it was hinted that using protein precipitation for ASOs, the recovery was increased to 25% when TEA was present in the extraction mixture. To date, there is no systematic report of protein precipitation method for ONTs, with and without the use of amines, achieving high recovery. In this study we present a simple and effective method that enables high recovery for multiple classes of ONTs from bi ological matrices through protein precipitation, assisted by am monia or other volatile amines, such as triethylamine (TEA) and diisopropylethylamine (DIPEA).

Given that traditional protein precipitation techniquesutilizing organic solvents result in low recoveries, we refer to this alternative approach as “enhanced protein precipitation” (EPP), which leads to a substantial improvement in ONT recovery post-protein precipitation. Here, the initial establishment of the EPP workflow is presented followed by a brief optimization of some key parameters. Finally, we demonstrate a proof-of-concept applicability of EPP for bioanalysis of oligonucleotides in various biological matrices such as mouse plasma and tissue.

## Experimental Section

Chemicals and Reagents. All ASOs and siRNA molecules were designed in-house and synthesized by Bio-Synthesis Inc. (Lewisville, TX). The type of chemistry and/or conjugation utilized in these compounds is detailed in Table 1. Their identity and purity were verified through liquid chromatography and mass spectrometry, while the reference quantities were determined by measuring the optical density at 260 nm. LC-MS grade methanol, acetonitrile, water, formic acid, and 1,1,1,3,3,3-Hexafluoro-2-propanol (HFIP) were procured from Fisher Scientific (Fisher Chemical, Fairlawn, NJ, USA). DNase/RNase-free water was obtained from Invitrogen (Austin, TX). LC-MS grade ammonia was sourced from Honeywell (Muskegon, MI), and N,N-Diisopropylethylamine (99.5%, biotech grade), ammonium formate, and other reagents, were purchased from Sigma-Aldrich (St. Louis, MO). CD-I mouse plasma (K2EDTA), gender-pooled, unfiltered, along with blank tissues, were acquired from Bio-IVT (Westbury, NY).

**TABLE 1.** ASO & siRNA Analytes and Internal Standards

| Name | MW (kDa) | Chemistry<sup>a</sup> | Use |
| --- | --- | --- | --- |
| ASO1 | 4.1 | MOE/DNA gapmer with PS backbone | Analyte |
| ASO2 | 6.4 | MOE/DNA gapmer with PS backbone | Analyte |
| ASO3 | 3.8 | MOE/DNA gapmer with PS backbone | Analyte |
| ASO4 | 7.1 | 5-10-5 MOE Gapmer with PS backbone | Internal Standard |
| siRNA1 | S: 13.3<br>AS: 7.4 | GalNAc Conjugated siRNA, mixed backbone with OMe and F modifications | Analyte |
| siRNA2 | S: 5.7<br>AS: 6.6 | Lipids Conjugated siRNA, mixed backbone with OMe and F modifications | Analyte |
| siRNA3 | S: 6.3<br>AS: 7.4 | Lipids Conjugated siRNA, mixed backbone with OMe and F modifications | Analyte |
| siRNA4 | S: 13.9<br>AS: 8.0 | GalNAc Conjugated siRNA, mixed backbone with OMe and F modifications | Internal Standard |

<sup>a</sup>aDNA, deoxyribonucleotides; MOE, 2'-O-(2-methoxyethyl)-oligoribonucleotides; OMe: 2'-O-methylated-oligoribonucleotides; F, 2'-Fluoro-oligoribonucleotides; PS, phosphorothioate linkage; GalNAc, N-Acetyl galactosamine, S: Sense strand, AS: antisense strand

Solutions and Standard Preparation. The EPP extraction solution is comprised of 1:1 (v/v) mixture of acetonitrile and methanol containing 1% ammonia (w/v). In cases where a reagent other than ammonia is evaluated, this is explicitly stated, and concentrations of the alternative reagent are adjusted to be equimolar to 1% (w/v) ammonia (approximately 600 mM) or other values justified for given experiments. For quantitative analyses, an internal standard (IS) was incorporated at a concentration of 100 ng/mL into the EPP extraction solution. Analyte reference materials were individually reconstituted in DNase/RNase-free distilled water to a final concentration of l0mg/mL. For qualitative, mechanistic, and recovery studies, working solutions of ASO1, ASO2, ASO3, siRNA1, and siRNA2 were spiked into plasma or other media to achieve a final concentration of 1000 ng/mL.Quantitative assessment. The MS/MS-based quantitative evaluation of plasma samples was conducted for ASO2, utilizing ASO4 as an internal standard (IS), with calibration standards prepared at concentrations of 1.00, 2.00, 5.00, 10.0, 20.0, 50.0, 100, 200, 500, 1000, and 2000 ng/mL. The lower limit of quantitation (LLOQ) and quality controls (QCs) were prepared at 1.00 ng/mL (LLOQ), 3.00 ng/mL (LQC), 750 ng/mL (MQC), and 1500 ng/mL (HQC) in plasma. HRMS-based quantitation of plasma samples was performed for siRNA3, with siRNA4 serving as the IS. Calibration standards in plasma were prepared at concentrations of 5.00, 10.0, 20.0, 50.0, 100, 200, 500, 1000, 2000, and 5000 ng/mL.

The LQC, MQC, and HQC for siRNA3 were prepared at 15.0, 750, and 3500 ng/mL, respectively, in plasma. Additionally, HRMS quantitation for siRNA3 was conducted on liver homogenized 1:9 (w/v) in blank plasma. Calibration standards were preparedat 50, 100, 200, 500, 1000, 2000, 5000, 10000, 20000, and50000 ng/ g tissue (5.00 to 5000 ng/ mL Homogenate) with 3 QC levels at 750 ng/ g tissue (LQC), 15000 ng/ g tissue, and 35000 ng/ g tissue. Cross-matrix quantitation of kidney and heart homogenates vs calibration curve made in liver homogenate was demonstrated at LQC level for each tissue type.

Sample Extraction. In this report, the extraction solvent-to-sample ratio is maintained at 4:1 (v/v) unless indicated otherwise (see Results and Discussions). Typically, a 100 μL aliquot of plasma or liver homogenate (called IX volume) is extracted with 400 μ1L of the pre-chilled EPP extraction solution (4X volume). The mixture is vigorously agitated for a minimum of 5 minutes at 2500rpm and subsequently centrifuged at 18,000 rcf for at least 20 minutes at 4°C. The supernatant is then collected and evaporated under a nitrogen flow using a plate-based evaporator (SPE Dry™ Biotage, Salem, NH). Following evaporation, the lyophilized sample resuspended in 100 μL of 10% methanol (aq) followed by a 60sec vortex at 2500 rpm. The reconstituted sample is subjected to an additional centrifugation step for 5 minutes at 18,000 rcf, with 90 μL of the clarified supernatant transferred to injection plate. 5 μL final sample was analyzed by ultra-high-performance liquid chromatography (UHPLC) coupled with either a high-resolution mass spectrometer (HRMS) operated in full scan mode or a triple quadrupole mass spectrometer (MS/MS). Additional details regarding EPP extraction using other reagents can be found in the Supporting Information.

UHPLC-HRMS and MS/MS. Chromatographic separation of ONTs was achieved using reversed-phase ion-pairing chromatography. A 2.1 mm x 50 mm Waters Premier BEH CI8 Oligo-nucleotide column was used. A binary mobile phase system was used including mobile phase A consisted of 10% methanol in water, and mobile phase B consisted of 70% methanol and 20 % acetonitrile in water, both mobile phases supplemented with 1% (v/v) 1,1,1,3,3,3-Hexafluoro-2-propanol (HFIP), 0.25% N,N-Diisopropylethylamine (DIPEA), and 2.5 μM ethylenediaminetetraacetic acid (EDTA). The flow rate of the mobile phase was set to 0.3 mL/min at a temperature of 75 °C. Further information is available in the Supporting Information. For mecha nistic studies, HRMS-based analyses were performed using either a Q-Exactive HF or an Orbitrap Exploris 120 (both Thermo Fisher Scientific, San Jose, CA), both instruments equipped with an H-ESI source operating in negative ion mode with a scan range of 650-2500m/z. MS/MS-based quantitation of ASO2 was conducted using the QTRAP 6500+ (Sciex, Framingham, MA) with an electrospray ionization (ESI) source operating in negative ion mode and employing multiple reaction monitoring (MRM) optimized for ASO2 and its internal standard (IS), ASO4 (Supporting Table S.1). HRMS-based quantitation of siRNA3 was executed using a Q-Exactive HF mass spectrometer operated in full scan mode. Additional details regarding these methodologies are provided in the Supporting Information.

Recovery calculations and data processing. In all cases recovery is presented as a ratio of chromatographic peak area for a given compound in extracted sample versus that in neat aqueous (Aq) solution at same concentration. All recovery measurements are performed as minimum duplicate measurements. Other data processing was done with the corresponding software from each MS vendor (Supporting Information). Graphs were plotted with GraphPad™ Prism 10 software.

## Results and Discussions

Establishing EPP. Development of a successful extraction method for ONTs via protein precipitation necessitates recognition of two distinct factors: (1) When biological matrix is absent, such as in an aqueous solution, the addition of a 4X organic solvent results in chromatographic peak splitting rather than precipitation, in addition to lowering the signal due to dilution; (2) Conversely, in the presence of a biological matrix, ONTs undergo actual precipitation upon the introduction of an organic solvent. Figure 2A presents the combined extracted ion chromatogram (combined XIC, explained in Supporting Information) of ASO1 in a neat Aq solution, serving as a reference signal. As depicted in Figure 2B, when a 4X 1:1 acetonitrile:methanol mixture (termed organic solvent) is added to a 100 μL aqueous solution of ONTs, resulting in an 80% organic content, a notable peak splitting of ASO1 during the IPRP chromatography is observed. This process results in elution of the majority of ASO 1 at the void volume (0.5 min) and a minor peak at the expected retention time (RT) of 2.7 min. Mass spectrometry confirmed that the early eluting peak at 0.4 minutes (z=-5 charge state) was indeed ASO1, and not due to crosstalk with background signals.

**Figure 1.**
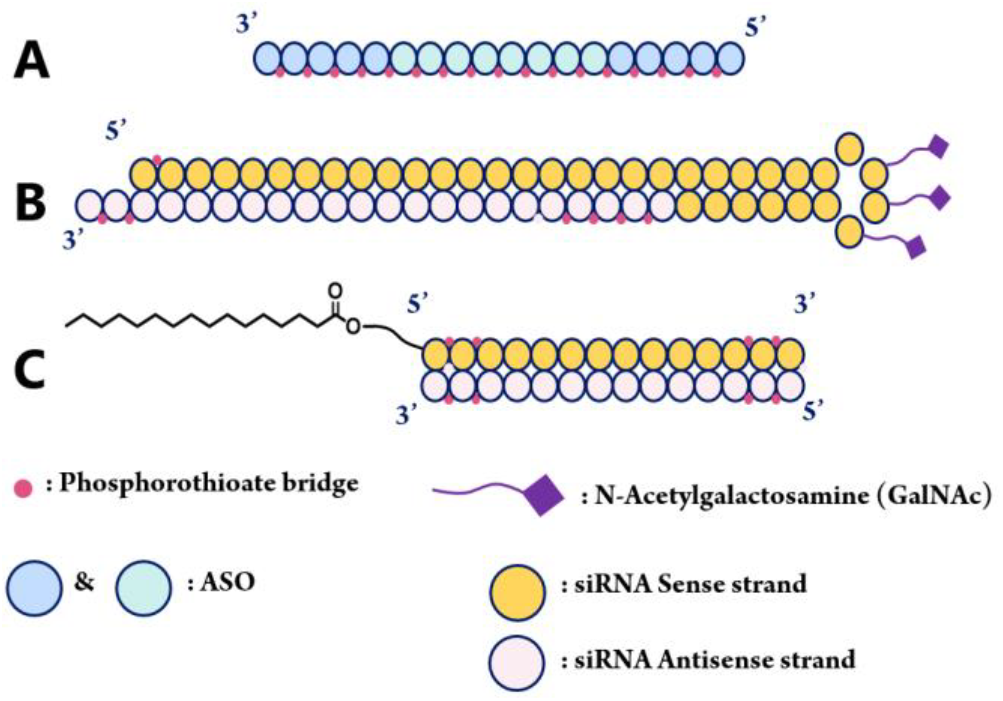
Representative structures for (A) ASO, (B) GalNAc conjugated siRNA, and (C) lipid conjugated siRNA

**Figure 2.**
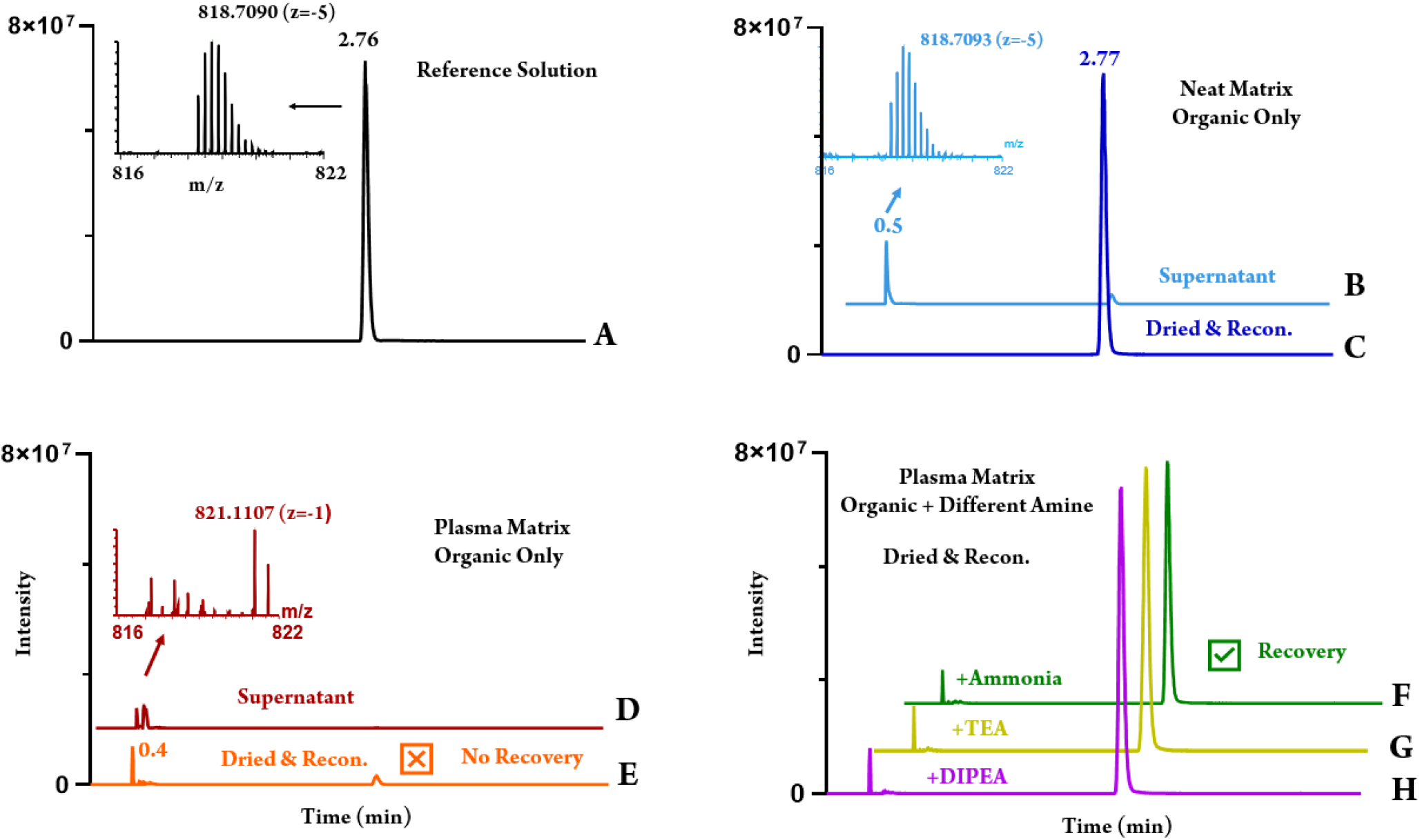
Combine Extracted Ion Chromatograms (XICs) of ASOl by combining XIC of all charge states ranging from 650-2500 m/z. (A) ASO1 peak at RT 2.6 mins in neat aqueous solution, (B) ASO1 peak post organic only EPP extraction in water observed as retaining in the void volume due to presence of 80% organic in the injection, (C) ASOl peak observed at the expected RT of 2.6mins post drying and reconstitution in 10% MeOH, (D) No ASO1 peak in void volume post organic only EPP extraction in plasma with 80% organic in the injection, (E) No ASO1 peak at expected RT post organic only EPP extraction in plasma with sample drying and reconstitution, (F) ASO1 peak at expected RT of 2.6mins post organic with ammonia EPP in plasma with sample drying and reconstitution, (G) ASO1 peak at expected RT of 2.6mins post organic with TEA EPP in plasma with sample drying and reconstitution, (H) ASOl peak at expected RT of 2.6mins post organic with DIPEAEPP in plasma with sample drying and reconstitution

Common strategies of peak shape improvement such as lowering the injection volume or increasing Aq content did not fully resolve this peak splitting, while it would decrease the sensitivity. If the typical practice of diverting void eluents to waste is followed, the unretained portion of ASO1 would not be detected, potentially leading to the erroneous conclusion of “precipitation.” This interpretation was confirmed bydrying the supernatant and reconstituting it to the original 100 μL volume using 10% methanolin water (termed reconcentration), where ASO1 nearly fully recovered at 90% at the anticipated retention time of 2.7 min (Figure 2C). In contrast, ONTs did truly precipitate within biological matrices upon the addition of organic solvent. When 100 μL plasma solutions of ASO1 were treated with 4X organic solvent, negligible recovery was observed both before (Figure 2D) and after (Figure 2E) reconcentration. Comparisons between these and the findings when matrix was neat Aq solution suggests that ONT loss through “precipitation” in biological samples is significantly associated with the presence of biological matrix components, likely proteins, rather than an inherent insolubility in solutions at 80% (v/v) organic content.

Incorporating ammonia or other small volatile amines in the extraction solution enhanced the recovery of ONTs from biological matrices. Figure 2F illustrates that the inclusion of 1% (w/v) ammonia (approximately 590 mM) in this organic solution (termed EPP extraction solution) facilitated the recovery of ASO1 from mouse plasma. Similarly enhanced recoveries were observed with the same molar concentrations of other small amines in extraction solution, such as triethylamine (TEA) (Figure 2G) and diisopropylethylamine (DIPEA) (Figure 2H), both achieving recoveries exceeding 90%. These trends were consistent across ASO2, ASO3, and various siRNAs, as described in subsequent discussions. Additional small volatile amines, including trimethylamine (TMA), tripropylamine (TPA), and hexylamine (HA), were also evaluated with satisfactory outcomes (Figure S.1). Ammonia was ultimately chosen as the EPP reagent due to its lower boiling point, a more manageable odor than certain alternatives like TMA, and better solubility than other amines. Other qualitative observations further justified the selection of ammonia (data not shown), for example that DIPEA resulted in reduced pellet thickness and less clean extracts, or tributyl amine (TBA) forming immiscible droplets.

Recovery does not depend on the order in which ammonia and organic solvent are added to the sample. Adding ammonia either before or after the organic solvent yielded comparable results with the addition of a combined solution, highlighted in Figure 3. Furthermore, when a solid pellet obtained post-centrifugation of a plasma sample exposed to 4X volume of organic solvent, where low recovery in supernatant is already established, was extracted by EPP the ONTs were recovered at near 100% recovery. These observations suggest that ammonia might act as disruptor of the interactions between ONTs and matrix, at any step of the precipitation and “rescue” ONTs from the matrix and keep them soluble in the supernatant. It is worth pointing out that although extraction from precipitated pellet involves additional steps than direct EPP, since biological samples are often limited, this approach potentially offers parallel extraction of molecules of different classes from the same biological sample (i.e. small molecule therapeutics/endogenous metabolite/lipid biomarkers) in supernatant of the organic extraction step, and ONTs from its pellet. This would be beneficial for studies where such comprehensive analysis is needed while it is not as easily achieved in other ONT extraction strategies such as SPE or Hybridization.

**Figure 3.**
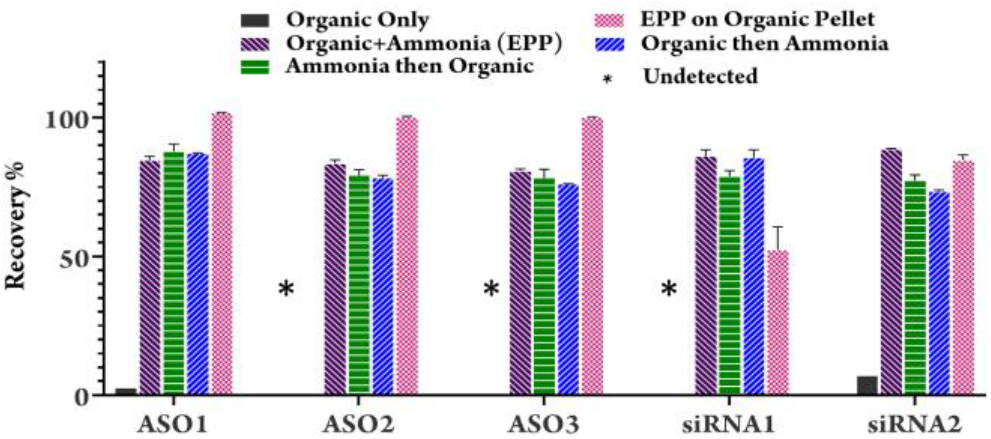
Effect of order of ammonia addition to different steps along the EPP extraction process to demonstrate no impact on recoveries for all 5 analytes. Furthermore, highlighting the role of ammonia in rescuing ONT from organic pellet (dotted bar plot)

EPP extraction for both strands of siRNAs were effective. EPP allows recovery of both sense and antisense strands from siRNAs from mouse plasma (Supporting Figure S2).

Ammonia Optimization. The effect of ammonia concentration in the extraction solution was examined in mouse plasma (Figure 4A). An optimal 1% (w/v) ammonia enhanced recovery for ASOs. For both siRNAs, addition of even 0.1% (w/v) ammonia increased the recovery to 70% which after addition of 1% (w/v) was further increased to above 80%. ASOs have higher phosphorothioate (PS) content in the backbone instead of phosphate (PO). Due to high PS content, ASOs are known to bind tightly to plasma proteins.^49^ This agrees with higher un-‘bound fractions observed for siRNAs^50^ than those reported for ASOs.^51^ As a result the hypothesis that excess ammonia interrupts protein-ONT interactions is strengthened,^47^ and the observation that ASOs require a higher % of ammonia compared to siRNAs to reach similar recovery corroborates that. Elevated concentrations of ammonia above 2% (w/v) resulted in formation of less desirable gel-like samples, and lack of clear supernatants after centrifugation, impeding further processing. For universality of future experiments, we chose 1% (w/v) ammonia to be included in EPP extraction solution.

**Figure 4.**
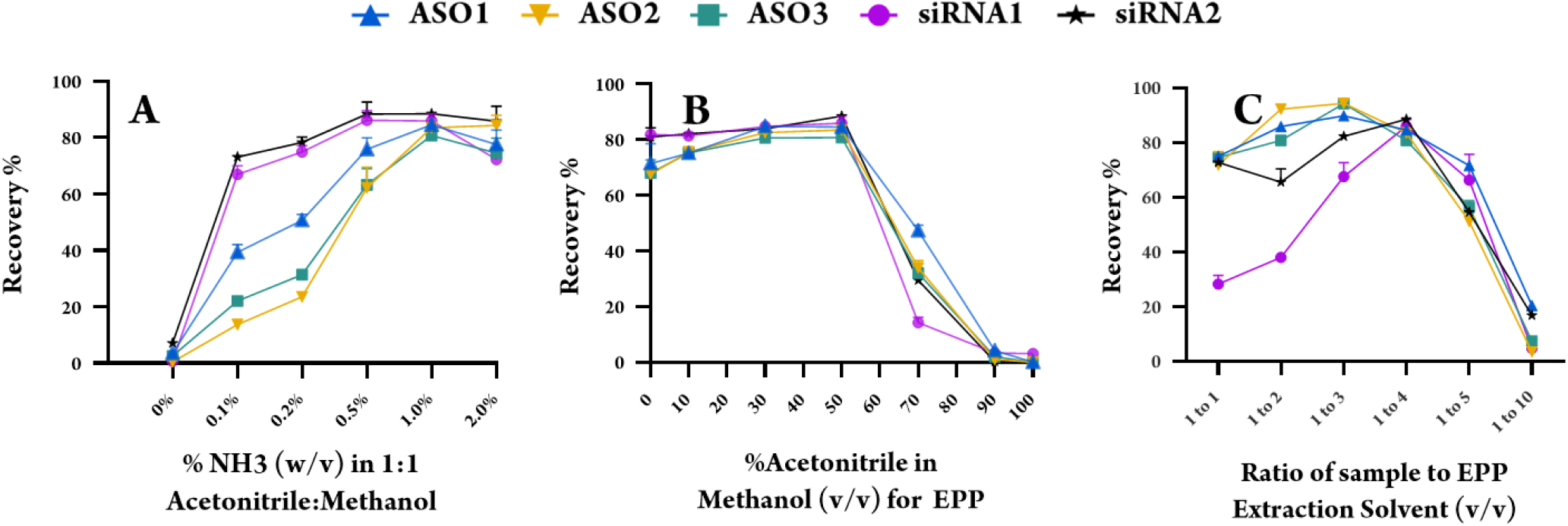
Optimization of extraction conditions. (a) Assessment of extraction recoveries for all 5 analytes with increasing ammonia concentration in EPP extraction solution (Recovery% calculated as peak area ratio of combined analyteXIC pre-spiked samples/combined analyte XIC in neat solution) (B) Assessment of composition of organic solvent for extraction solution. (C) Assessing Solvent to-Sample ratio by evaluating recoveries of all 5 analytes in plasma by EPP extraction using organic with ammonia

Evaluation of Organic Solvents. Recovery of all ONTs from mouse plasma seem to be sensitive to the amount of acetonitrile in the extraction solution. As shown in Figure 4B, when extraction solutions contained above 50% acetonitrile recovery dropped significantly. While having higher than 50% methanol in the extraction solution showed reasonable recovery across different ONTs, it muddled supernatant clarity and introduced background noise, eventually impacting column integrity. The dried solid residues after extraction with 50%-100% acetonitrile and 1% (w/v) ammonia were visibly cleaner and lesser than those obtained from high methanol content which corroborates the negative impact observed on chromatography column after sustained exposure to extracts obtained from methanol including 1% (w/v) ammonia. 50% acetonitrilein methanol was identified as an optimum organic composition resulting in acceptable recovery resulting in clean samples for prolonged LCMS analyses.

Solvent-to-Sample Ratio Optimization. Different EPP extraction solvent volumes were tested while the final reconstitution solvent and volume remained the same for all samples (Figure 4C). Excessive solvent (10X) diminished recovery, denoting a true precipitation of ONTs at very high organic contents. Less than 4X volumes resulted in a similar recovery although caused elevated column back pressure and more ion suppression during LCMS analysis. Opting for 4X volume avoided these issues. Table 2 summarizes the optimized extraction parameters for EPP experiment.

**TABLE 2.** Optimized EPP extraction conditions

| Parameter | Suggested value for ASOs | Suggested value for siRNAs |
| --- | --- | --- |
| %NH<sub>3</sub> in the extraction solution (w/v) | 1.0% | 0.5% - 1.0% |
| Ratio of Extraction Solution to Sample | 3:1 - 4:1 | 4:1 |
| % Acetonitrile in Methanol | 30%-50% | 30%-50% |

Recovery of different ONTs with EPP and comparison with SPE. Table 3 shows recoveries obtained for different ONTs using EPP. We observed that for ASOs, EPP offers similar recovery compared to SPE while for siRNA it results in improved recovery. For instance, recovery for lipid-conjugated siRNAs (above 80%) where SPE lagged under 50%. Other ONT classes performed similarly across EPP and SPE. Summarized qualitative observations and matrix interference for each extraction are detailed in Supporting Information. (Supporting Figure S.3).

**TABLE 3.** Precision and Accuracy of ASO2 Quantitation in mouse plasma

| Extraction Method | ASO1 | ASO2 | ASO3 | siRNA 1 |  | siRNA 2 |  |
| --- | --- | --- | --- | --- | --- | --- | --- |
|  |  |  |  | Antisense | Sense | Antisense | Sense |
| EPP | 99% | 96% | 93% | 86% | 89% | 85% | 84% |
| OligoWorks | 96% | 91% | 97% | 43% | 42% | 23% | 13% |
| Clarity SPE | 94% | 89% | 87% | 52% | 60% | 15% | 8% |

In the biological matrix, ONTs interact tightly with proteins via electrostatic pairing of phosphate backbone with positively charged residues like lysine and arginine or via weaker hydrogen bonds. In EPP, ammonia (roughly 600 mM) is 1000-fold in excess compared to the most abundant protein in mouse plasma (albumin, at 600 μM). Disruption of ONT-protein interaction achieves ∼80% recovery during, before, or after precipitation induced by organic solvent. Volatile amines serve this purpose by competing with protein cationic residues for interaction with negatively charged ONTs. Understanding how amines perform this requires further experiments. However, 3 other notable observations may provide a starting point for future research: (1) lack of recovery when weak acids were used in extraction solution (data not shown), (2) partial recovery when extraction solution contained small amounts of strong base like sodium hydroxide,(Supporting Figure S.4), (3) partial recovery when sample was pre-incubated with large amount of soluble ammonium formate at same molarity as 4X volume of 1% ammonia would provide; roughly 2.4 M (Supporting Figure S.4). In addition to ionic interactions, amines and their ammonium ions can interact with nucleotides through hydrogen bonding with different moieties of ONTs.^38,5–55^ Lastly, ammonia in aqueous or organic solvent increases the pH, provides ionic and H-bondin-teractions all hypothesized to be desirable factors leading to rescue of ONTs from matrix proteins. A representation of such plausible interaction of amines with ONTs leading to their recovery from the biological matrix is shown in Supporting Figure S.5.

### Analytical Method Qualification

A preliminary qualification of the EPP extraction method for the bioanalysis of ONTs was conducted. Analytical method qualification for two categories of ONTs; ASOs and siRNAs, is presented here while a different MS detection strategy was used for each category to showcase compatibility of EPP with different MS platforms commonly used for small and large molecule bioanalysis. For ASOs, we conducted MS/MS-based quantitation in MRM mode on a triple quadrupole mass spectrometer, while for siRNAs we chose full scan HRMS for the analysis. In each case calibration curves and associated quality control (QC) samples were analyzed and their accuracy and precision values were assessed. For ASO2 in mouse plasma, the precision and accuracy of QC samples were assessed through inter-day and intra-day analyses, utilizing two calibration curves and four QC levels, each with six replicates. For siRNA3, assessment was conducted in mouse plasma and liver homogenates, each with two calibration curves and 3 QC levels within a single-day evaluation. Additionally, since other tissue types are often limited, cross-tissue quantitation of siRNA3 in mouse kidney and heart homogenates was examined via use of liver homogenate calibration curve. Despite EPP compatibility with automation, all across three distinct spike-in samples, calibration curves, and QC samples were man ually prepared.

Analytical Method Qualification for ASO2. The precision and accuracy of ASO2 QC samples are documented in Table 4, using ASO4 as the internal standard (IS). MS/MS parameters for ASO2 and ASO4 are outlined in Table S.1 (Supporting Information). For both ASO2 and the IS, the phosphorothioate fragmention corresponding to m/z 95.0 was monitored. The response factor, expressed as the ratio of ASO2 to ASO4 peak areas, was plotted against the concentration of ASO2 utilizing linear regression with 1/x^2^ weighting, covering a range from 1 ng/mL to 2000 ng/mL (2000x). This methodology achieved linearity of 0.9972, 0.9929, and 0.9914 across three distinct runs. On-column carryover caused by upper limit of quantitation (ULOQ) samples in the calibration curve (2000 ng/mL) presented a notable challenge. This issue was effectively mitigated by injecting four matrix blanks, which reduced carryover signal to 20% of the lower limit of quantitation (LLOQ, 1 ng/mL). Figure 5 illustrates representative extracted ion chromatograms (XICs) for ASO2 and the IS in blank mouse plasma (Figure 5A), as well as at LLOQ(Figure 5B) and ULOQ levels (Figure 5C) accompanied by stability of IS across the run (Figure 5D) an ASO2 calibration curve in mouse plasma (Figure 5E). Although IS crosstalk was noted in the analyte channel (Figure 5B), the opposite crosstalk was not observed. Due to the chromatographic resolution achieved, this crosstalk did not impede accurate quantification.

**TABLE 4.** Precision and Accuracy of ASO2 Quantitation in mouse plasma

| QCs in Plasma |  |  |  |  |
| --- | --- | --- | --- | --- |
| no | LLOQ 1 | QCL 3 | QCM 750 | QCH 1500 |
|  | measured conc<br>(ng/mL) | measured conc<br>(ng/mL) | measured conc<br>(ng/mL) | measured conc<br>(ng/mL) |
| 1 | 1.140 | 2.66 | 840.0 | 1610 |
| 2 | 1.170 | 2.75 | 785.0 | 1590 |
| 3 | 1.220 | 3.45 | 853.0 | 1550 |
| 4 | 1.150 | 2.96 | 837.0 | 1660 |
| 5 | 1.050 | 2.72 | 863.0 | 1540 |
| 6 | 0.851 | 2.91 | 764.0 | 1580 |
| n | 6 | 6 | 6 | 6 |
| mean | 1.097 | 2.91 | 823.7 | 1588 |
| SD (n-1) | 0.13 | 0.29 | 39.77 | 43.55 |
| CV% | 12.1% | 9.9% | 4.8% | 2.7% |
| accuracy% | 109.7% | 96.9% | 109.8% | 105.9% |
| n | 18 | 18 | 18 | 18 |
| mean | 1.055 | 2.85 | 778.6 | 1519 |
| SD (n-1) | 0.15 | 0.19 | 73.62 | 126.14 |
| CV% | 16.6% | 6.7% | 10.3% | 9.1% |
| accuracy% | 105.5% | 95.0% | 103.8% | 101.3% |

**Figure 5.**
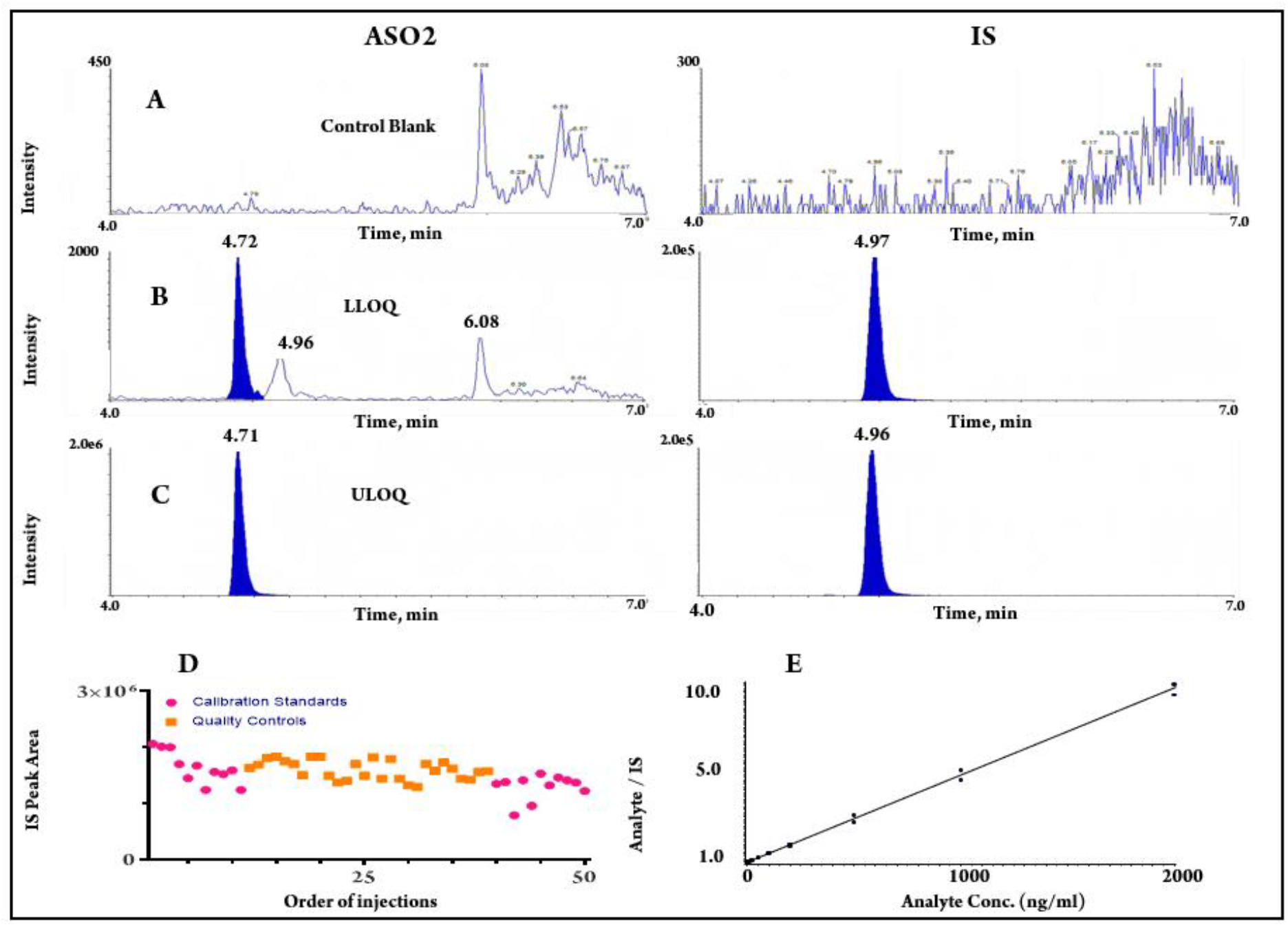
LC-MS/MS chromatograms of ASO2 and the IS from (A) blank mouse plasma sample, (B) spiked LLOQ sample at lng/mL in mouse plasma, (C) spiked ULOQ sample at 2000ng/mL in mouse plasma, (D) IS peak area response over the course of data acquisition (E) Calibration curve of ASO2 from Ing/mL - 2000ng/mL in mouse plasma.

Analytical Method Qualification for siRNA3. The evaluation of siRNA3 was conducted using HRMS detection following EPP extraction from mouse plasma and tissue homogenate samples. While the focus was on the antisense strand, it is noteworthy that HRMS detection allows for potential detection of both antisense and sense strands and their potential metabolites. Liver samples were homogenized in blank plasma, as detailed in the Methods section.

Figure 6 shows the combined XICs for siRNA3 and its corresponding IS in blank mouse plasma (Figure 6A), at LLOQ(Figure 6B), and ULOQ levels (Figure 6C), and the calibration curve for siRNA3 in both mouse plasma (Figure 6D) and liver homogenate (Figure 6E). IS crosstalk was observed in the analyte channel (Figure 6B) however this posed no challenge due to chromatographic resolution of the peaks. Using HRMS for siRNAs, in range of 5 to 5,000 ng/mL a linearity of 0.998 was achieved for plasma samples and 0.997 for liver homogenates between 5 to 5,000 ng/mL_homogenate_ (50 to 50,000 ng/g_issue_). Table 5 summarized the analytical qualification outcomes for siRNA3 in mouse plasma and liver homogenate after extraction with EPP. Cross tissue quantitation was successful for kidney and heart tissues.

**TABLE 5.**
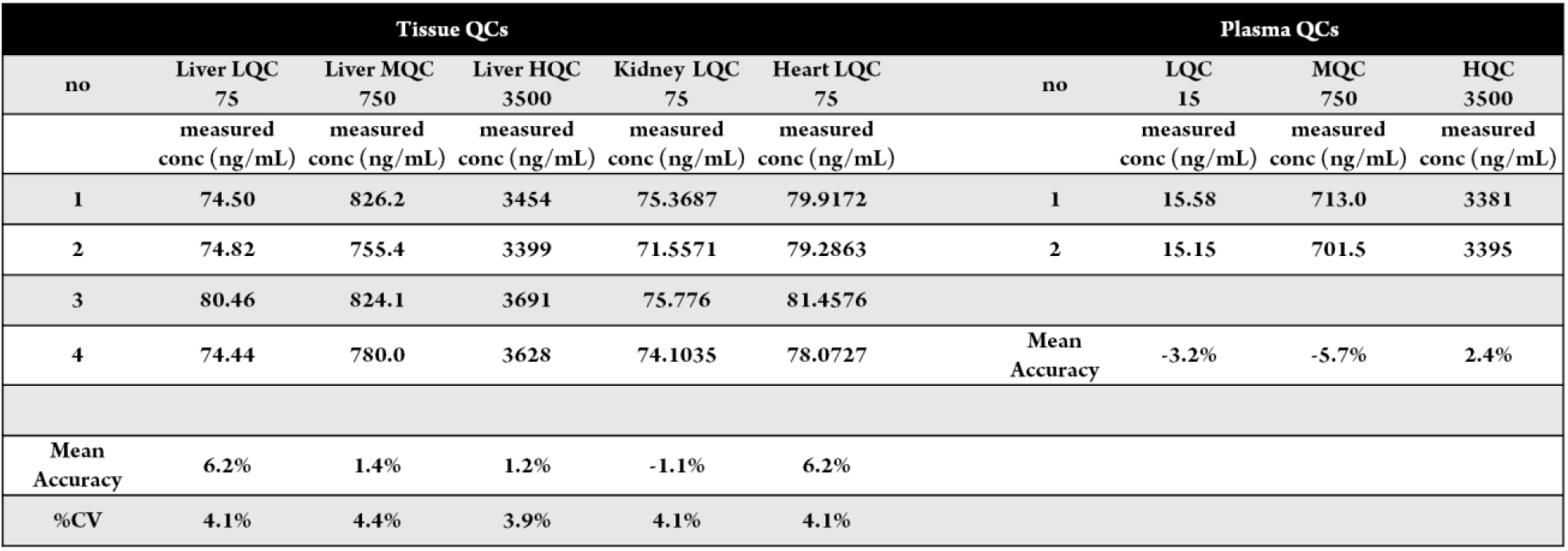
Analytical method qualification for siRNA 3 with mouse plasma and mouse liver QC samples quantitated against calibra tion curves in matching matrices

| Tissue QCs |  |  |  |  |  | Plasma QCs |  |  |  |
| --- | --- | --- | --- | --- | --- | --- | --- | --- | --- |
| no | Liver LQC<br>75 | Liver MQC<br>750 | Liver HQC<br>3500 | Kidney LQC<br>75 | Heart LQC<br>75 | no | LQC<br>15 | MQC<br>750 | HQC<br>3500 |
|  | measured<br>conc (ng/mL) | measured<br>conc (ng/mL) | measured<br>conc (ng/mL) | measured<br>conc (ng/mL) | measured<br>conc (ng/mL) |  | measured<br>conc (ng/mL) | measured<br>conc (ng/mL) | measured<br>conc (ng/mL) |
| 1 | 74.50 | 826.2 | 3454 | 75.3687 | 79.9172 | 1 | 15.58 | 713.0 | 3381 |
| 2 | 74.82 | 755.4 | 3399 | 71.5571 | 79.2863 | 2 | 15.15 | 701.5 | 3395 |
| 3 | 80.46 | 824.1 | 3691 | 75.776 | 81.4576 |  |  |  |  |
| 4 | 74.44 | 780.0 | 3628 | 74.1035 | 78.0727 | Mean<br>Accuracy | -3.2% | -5.7% | 2.4% |
| Mean<br>Accuracy | 6.2% | 1.4% | 1.2% | -1.1% | 6.2% |  |  |  |  |
| %CV | 4.1% | 4.4% | 3.9% | 4.1% | 4.1% |  |  |  |  |

**Figure 6.**
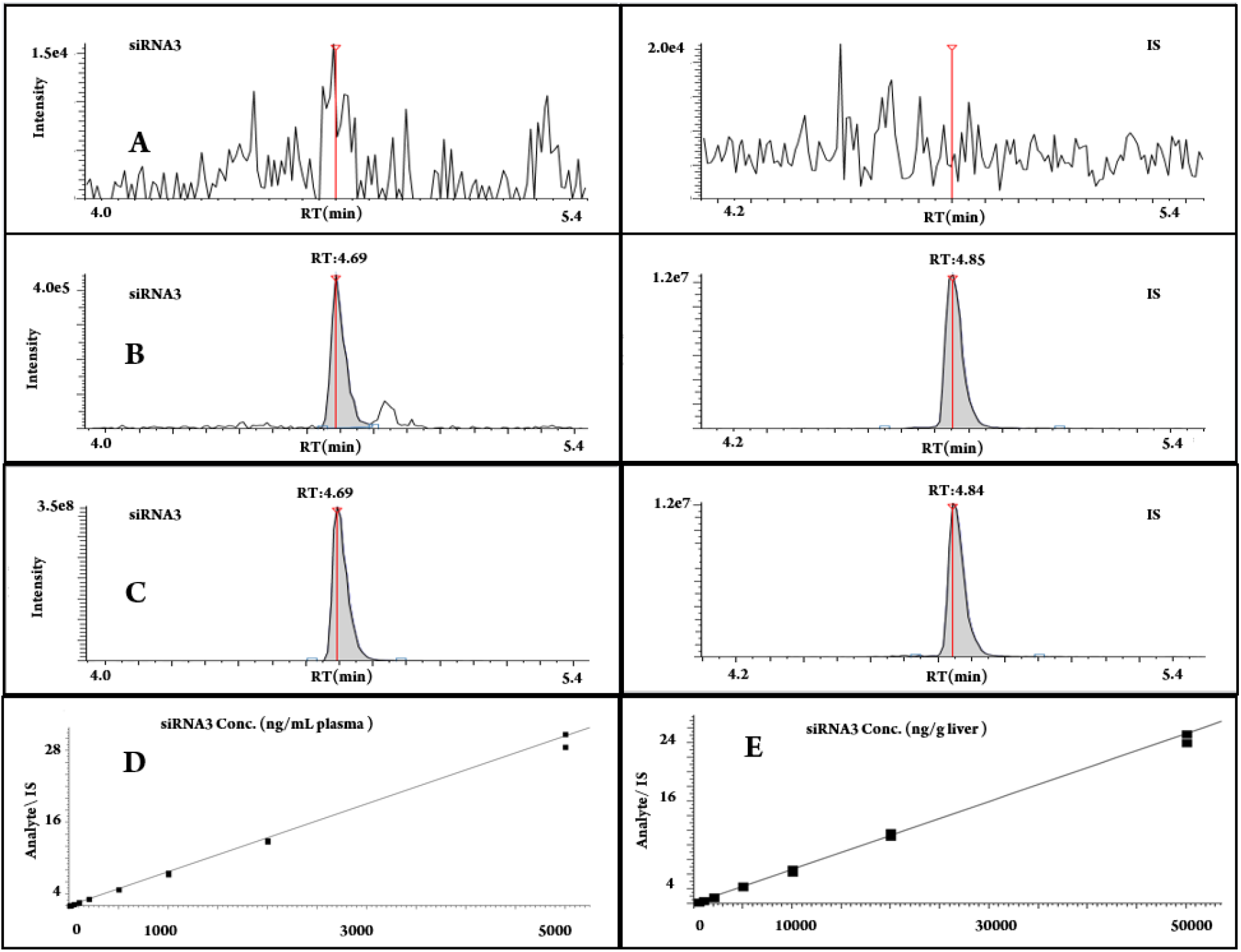
Combined Extraction Ion Chromatograms (XICs) of siRNA3 and the IS from (A) blank mouse plasma sample, (B) spiked LLOQ sample at 5ng/mL in mouse plasma, (C) spiked ULOQ sample at 5000ng/mL in mouse plasma, (D) Calibration curve of siRNA 3 from 5ng/mL - 5000 ng/mL in mouse plasma, (E) Calibration curve of siRNA 3 from 5ng/mL - 5000 ng/mL (equivalent to 50- 50,000 ng/g tissue) in mouse liver homogenate.

Interestingly, the calculated molar equivalents of the obtained LLOQs were approximately 0.15 nM for ASO2 and 0.13 nM for siRNA3, underscoring that similar molar sensitivity achieved for EPP across various compounds and across differ ent MS workflows for quantitation.

## Conclusions

In this study we developed and qualified enhanced protein precipitation (EPP), an effective and practical method for the bioanalysis of oligonucleotides which overcomes many limitations of existing approaches, including high cost, lengthy assay time, and intensive method development. EPP is protein precipitation with the addition of ammonia or other low molecular weight volatile amines such as TEA or DIPEA. This unique optimization in the standard precipitation procedure boosts recovery of ONTs when extracted from biological matrices enabling a bioanalytical extraction method that is rapid, cost-effective, and scalable. EPP is applicable for chemically diverse ONTs, including ASOs and siRNAs and associated novel chemical conjugates of these molecules. We report that ammonia is ideally suited as an additive for this application due to several practical advantages, including ease of evaporation, cost, and post-precipitation extract consistency. We speculate that the enhancement of ONT recovery by amines is via disruption of interactions between proteins and ONTs, leading to their res cue from matrix during the extraction step. While some properties of ammonia like hydrogen bond donor/acceptor activity as well as high pH may have other critical roles in the success of EPP; more evidence on this matter is needed and will be evaluated in future studies. Finally, we evaluated the applicability of EPP for the bioanalysis of biological samples, demonstrating excellent limits-of-quantitation, linear dynamic range, and good reproducibility.We anticipate that EPP will become a core extraction method for oligonucleotide bioanalysis, facilitating ONTs drug discovery and development.

## ASSOCIATED CONTENT

### Supporting Information

The Supporting Information is available free of charge. Detailed experimental section, data processing, recovery when other amines used, comparison of recovery for sense and antisense strands for siRNAs, extraction with inorganic salts and sodium hydroxide, MRM optimized parameters for ASO2 and IS, and schematic view of proposed interactions between proteins and ONTs as well as amines and ONTs (PDF)

## ACKNOWLEDGMENT

Authors wish to acknowledge Dr. Xiao-Yi Xiao, Dr. Jianzhong Chen, Dr. Maryam Yahyaee Anzahaee, Jose Paredes, Gargi Kasture, Dr. Kartik Temburnikar, Dr. Yogesh Shelke and Professor Daniel Jones for the scientific discussions and data review.

