## Supplementary Data for "Enhanced protein precipitation with ammonia enables rapid, universal extraction of oligonucleotides for bioanalysis"

### METHODS:

#### Solutions and Standard Preparation

Individual stock solutions of ASOs and siRNAs were prepared by dissolving dry solid in DNAase/RNAse free water at 10 mg/mL. Working solutions were prepared as a mixture of the ASO1, ASO2, ASO3, siRNA1, and siRNA2 (called ONT mixture) at 100,000 and 10,000 ng/mL by serial dilutions from the stock solution in water. All stock and working solutions were stored at 4°C when not in use.

#### Neat and Plasma sample preparation:

10 uL of the 10,000 ng/mL solution was spiked in 90 uL of mouse plasma to yield 100 uL of sample at 1,000 ng/mL. For neat solutions 10 uL of the 10,000 ng/mL solution was spiked in 90 uL of DNAs/RNAse free water to yield 100 uL of sample at 1,000 ng/mL.

#### Extraction solutions:

The organic-only extraction solution was prepared by mixing acetonitrile and methanol at 1 to 1 ratio (termed organic solvent). EPP extraction solution was made by spiking ammonia in organic solvent at 1% (w/v). For example, 0.4 mL of concentrated ammonium hydroxide solution, at 25% ammonia, was spiked in 9.6 mL of acetonitrile:methanol mixture to create 10 mL of extraction solution. Extraction solutions for other amines were made at 600 mM concentration in the organic solvent. Extraction solutions were pre-chilled in fridge prior to extraction.

#### Sample extraction:

A 100 uL sample (termed 1X volume) was extracted with 400 uL of the extraction solution (4X) in 0.75 mL Thermo matrix microtiter tube. The tube was capped tightly, and the mixture was vortexed vigorously at 2500 rpm for a minimum of 5 minutes. The mixture was then centrifuged at 18000 g for a minimum of 20 minutes. Supernatants were transferred into a separate microtiter tube and dried under nitrogen flow. Lyophilized samples were then reconstituted in 10% methanol in water, followed by vortexing at 2500 rpm and 5 minutes of centrifugation at 18000 g. 90 µL of supernatant was transferred to injection plates where 5 uL was injected to LCMS. All extractions were done as two technical replicates.

#### Establishing EPP:

For neat solutions: Two separate duplicate sets of 100 µL of neat solution with 1000 ng/mL ONT mixture were extracted with organic extraction solution. Supernatant of one set was injected into LCMS directly after vortex at 2500 rpm for 10 minutes and centrifugation at 20 minutes. Supernatant from another set was dried down and processed as mentioned above before injection to LCMS. For plasma samples: Duplicate 100 µL plasma samples with 1,000ng/mL of ONTs were extracted with 400 µL of EPP extraction solution and separately with alternative EPP extraction solutions that contained 600 mM TEA and 600 mM DIPEA instead of ammonia. Samples were processed by vortexing, centrifuging, drying down and reconstitution before LCMS analysis as mentioned above. LCMS analysis was done with UHPLC-HRMS using LCMS Method 1 described below.

#### Evaluating the order of ammonia and organic solvent addition, and EPP extraction from organic pellet:

4 sets of 100 µL plasma samples with 1,000ng/mL of ONTs were prepared in duplicate. For the first set, 400 µL of organic solvent was added and after vortexing 16 µL of 25% ammonia was spiked. To the second set first 16 µL of 25% ammonia was added and after vortexing 400 µL of organic solvent was added. For the third set, regular EPP extraction was performed. These three sets were then vortexed more and processed as mentioned above before LCMS analysis. For the fourth set, only the organic solvent was added, vortexed and centrifuged, supernatant was removed, and 100 µL water was added and pellet was pulverized in the water solution followed by extraction with 400 µL of EPP solution and processing as mentioned above before LCMS analysis. Samples were analyzed with UHPLC-HRMS using LCMS Method 2 described below.

#### Optimization of Ammonia:

EPP extraction solution was made with 0.0%, 0.1%, 0.2%, 0.5%, 1%, and 2% (w/v) ammonia. Duplicate 100 µL plasma samples with 1,000ng/mL of ONTs were extracted with 400 µL of each of the extraction solutions. Samples were then processed as mentioned above followed by LCMS analysis. LCMS was done with UHPLC-HRMS using LCMS Method 1 described below.

**Evaluation of organic solvents:**

**Extraction solutions with 1% (w/v) ammonia were made in acetonitrile, methanol, and 1:1 mixture of the two. Duplicate 100 µL plasma samples with 1,000ng/mL of ONTs were extracted with 400 µL of each of the extraction solutions. Samples were then processed as mentioned above followed by LCMS analysis. ASO4 and siRNA4 at 400 ng/mL were added to the final extracts as post-spike to evaluate the matrix effect. Matrix effect was calculated by ratio of [peak area for post-spike/peak are for reference neat solution] represented as percentage. LCMS was done with UHPLC-HRMS using LCMS Method 2 described below.**

**Evaluation of sample to extraction solution volume ratio:**

**EPP extraction solution was prepared as mentioned above. Duplicate 100 µL plasma samples with 1,000ng/mL of ONTs were extracted with varying volumes of extraction solution at 100 µL, 200 µL, 300 µL, 400 µL, 500 µL, and 1000 µL to create 1:1, 1:2, 1:3, 1:4, 1:5, and 1:10 (v/v) ratio of sample to extraction solution. Mixtures were processed as mentioned above and analyzed by LCMS. LCMS was done with UHPLC-HRMS using LCMS Method 2 described below.**

**Extraction evaluation of lipid conjugated siRNA2 using Phenomenex Clarity OTX, Waters OligoWorks & EPP**

**3 sets of duplicate 100uL plasma samples with 1,000ng/mL of siRNA 2 were prepared. The first set of samples were extracted by following the EPP protocol described in the paper. The second and third set of samples were extracted via the standard Phenomenex Clarity OTX protocol and Waters OligoWorks protocol respectively. Phenomenex Clarity OTX protocol and Waters OligoWorks extraction method were followed as per manufacturer protocol as referenced below.**

**Phenomenex Clarity OTX:** https://www.phenomenex.com/products/clarityspe-sample-preparation-products/clarity-otx

**Waters OligoWorks:** https://www.waters.com/nextgen/us/en/library/application-notes/2024/extraction-of-oligonucleotides-from-plasma-samples-across-multiple-species-using-oligoworks-spe-microplate-kit.html

**Common chromatography parameters:**

**A 2.1 mm x 50 mm Waters Premier BEH C18 Oligonucleotide column was used in all experiments. Also gradient elution was used for all experiments with two mobile phases including mobile phase A (termed solvent A) consisted of 10% methanol in water, and mobile phase B (termed solvent B) consisted of 70% methanol and 20 % acetonitrile in water, both mobile phases supplemented with 1% (v/v) 1,1,1,3,3,3-Hexafluoro-2-propanol (HFIP), 0.25% N,N-Diisopropylethylamine (DIPEA), and 2.5 µM EDTA. The flow rate of the mobile phase was set to 0.3 mL/min at a temperature of 75 °C. 5 µL sample was injected for analysis.**

**LCMS Method 1: Separation was at flow rate of 0.3 mL/min and holding elution at 5% B for 0.1 min followed by a linear ramp to 10% B at 0.2 min, then to 20% B at 3 min, and then to 99% B at 5min keeping up to 8min and decreasing %B back to 5% at 8.1 min and holding for the next two minutes. HRMS analysis was done by a Thermo Orbitrap Exploris 120 equipped with H-ESI source operating in negative ionization mode with these conditions: -3000 volts, Sheath Gas 35, Aux Gas 15, and Sweep gas 0, Vaporizer Temperature of 320 $^0$C, RF lens 65, and Capillary Temperature 300 $^0$C. Scan range was 650 /z to 2500 m/z with 120,000 Resolution. Normalized AGC target of 100% and Automatic Maximum Injection Time. Combined extracted ion chromatograms (Combined XICs) for each of the ASOs and the antisense strands of each siRNA were created and processed by Thermo TraceFinder Version 5.2 and FreeStyle Version 1.8 software. Recovery was defined as the ratio of [peak area of analyte after extraction/peak area of analyte in neat solution] and represented as percentage.**

**LCMS method 2: Separation was done at flow rate of 0.3 mL/min and holding elution at 5% B for 0.2 minutes followed by a linear ramp to 25% B at 3 min, then to 99% B at 5 min, keeping up to 8 min, and decreasing %B back to 5% at 8.1 min and holding for the next two minutes. HRMS analysis was done was done by a Thermo QExactive HF machine equipped with H-ESI source operating in negative ionization mode with these conditions: -4000 volts, Sheath Gas 35, Aux Gas 12, and Sweep gas 0, Heater Temperature of 320 $^0$C, RF lens 65, and Capillary Temperature 300 $^0$C. Scan range was 650 /z to 2500 m/z with 240,000 Resolution. AGC target was 5e6 and the Maximum Injection Time was 250 ms. Combined extracted ion chromatograms (Combined XICs) for each of the ASOs and the antisense strands of each siRNA were created and processed by Thermo TraceFinder Version 5.2 and FreeStyle Version 1.8 software. Recovery was defined as the ratio of [peak area of analyte after extraction/peak area of analyte in neat solution] and represented as percentage.**

#### Analytical method qualification for ASO2

Standard solutions for ASO2 were prepared in mouse plasma in a 2000-fold range at concentrations of 1.00, 2.00, 5.00, 10.0, 20.0, 50.0, 100, 200, 500, 1,000, and 2,000 ng/mL. The lower limit of quantitation (LLOQ) and quality controls (QCs) were prepared at 1.00 ng/mL (LLOQ), 3.00 /mL (LQC), 750 ng/mL (MQC), and 1500 ng/mL (HQC) in mouse plasma. Duplicate calibration curve and 6 replicates of each QC were analyzed with LCMS/MS. ASO4 at 100 ng/mL was dissolved as IS in the EPP extraction solution. After extraction, 5 µL of each sample was separated by an Agilent 1290 UHPLC system and Sciex 6500+ triple quadrupole. Separation was at flow rate of 0.3 mL/min and holding elution at 5% B for 0.4 min followed by a linear ramp to 10% B at 0.5 min, then 15%B at 2.5 min, followed by 25%B at 5 min, and then 95%B at 6 min with holding at 95%B till 7 mins. Lastly, re-equilibrating the column at 5%B at 7.1 min for the next 1.5 min, thus finishing the gradient with a total 8.5 min run-time. MS/MS detection in multiple reaction monitoring (MRM) modes was done by monitoring the selected precursor and fragment ions shown in Supporting Table S.1 using the specified collision parameters.

Data processing was done by Sciex Analyst software 1.7. The response factor, expressed as the ratio of ASO2 to ASO4 peak areas, was plotted against the concentration of ASO2 utilizing linear regression with $1/x^2$ weighting, covering a range from 1 ng/mL to 2,000 ng/mL (2000x).

#### Analytical method qualification for siRNA3

For mouse plasma: pooled K2EDTA CD-1 mouse plasma from BioIVT (Westbury, NY) was used to prepare standard calibration samples. Calibration standards for siRNA3 in plasma were prepared at concentrations of 5.00, 10.0, 20.0, 50.0, 100, 200, 500, 1000, 2000, and 5000 ng/mL. The LQC, MQC, and HQC were prepared at 15.0, 750, and 3500 ng/mL, respectively, in plasma. Duplicate calibration curve and duplicate set of QCs were extracted with EPP extraction solution containing 100 ng/mL siRNA4 as IS. Samples were then analyzed by injecting 5 µL of final samples in a Thermo Vanquish F UHPLC followed by high resolution MS (HRMS) in full scan mode. Separation gradient was done by flow rate of 0.3 mL/min and holding elution at 5% B for 2 minutes followed by a linear ramp to 15% B at 2.1 min, then to 20% B at 5 min, 99% B at 8 min keeping it until 11 min, decreasing B to 5% at 11.1 min and keeping for 3 more minutes. HRMS analysis was done was done by a Thermo Q Exactive HF machine equipped with H-ESI source operating in negative ionization mode with these conditions: -4000 volts, Sheath Gas 35, Aux Gas 12, and Sweep gas 0, Heater Temperature of 320 $^0$C, RF lens 65, and Capillary Temperature 400 $^0$C. Scan range was 650 /z t o2500 m/z with 240,000 Resolution. AGC target was 3e6 and the Maximum Injection Time was 200 ms. Combined XIC for antisense strands were created and processed by Thermo TraceFinder Version 5.2 software. The response factor, expressed as the ratio of siRNA3 to siRNA4 peak areas, was plotted against the concentration of siRNA3 utilizing linear regression with $1/x^2$ weighting, covering a range from 5 ng/mL to 5000 ng/mL (1000x).

For Tissue: Blank mouse liver tissues from BioIVT (Westbury, NY) were weighed and homogenized in pooled K2EDTA CD-1 mouse plasma from BioIVT (Westbury, NY) at 1:9 (w/v) ratio to obtain a 100 mg/mL homogenate. BD FastPrep-24™ 5G homogenizer was used with 2 rounds of 20-seconds homogenization with 10 second interval. Over-night stability of the compounds in liver-in-plasma homogenates were confirmed (data not shown).Calibration standards were pre-pared at 50, 100, 200, 500, 1000, 2000, 5000, 10000, 20000, and 50000 ng/g tissue (5.00 to 5000 ng/mL Homogenate) with 3 QC levels at 750 ng/ g tissue (LQC), 15000 ng/ g tissue , and 35000 ng/ g tissue . Cross-matrix quantitation QC samples for kidney and heart were made in liver homogenate at LQC level for each tissue type. Duplicates of 100 µL calibration standards and 4 replicates of QC samples were extracted with EPP followed by same sample processing as plasma mentioned above. Analysis was done with the same LCMS method used for plasma.

#### Combine extraction ion chromatogram (XIC) for each compound in HRMS:

Combined XIC is made by combining individual m/z values for the 7 most abundant isotopes of all the predicted charge states of a given analyte that falls in the scan range (m/z 650 to m/z 2500). The said values for each compound were calculated using internal software based on an algorithm explained by Goraczko. A. J.[1]

**TABLES:**

**Supporting Table S.1. MRM Transitions and Parameters for ASO2 and ASO4 (IS)**

| ID | Q1 mass | Q3 mass | Dwell Time (ms) | DP (volts) | EP (volts) | CE (volts) | CXP (volts) |
|---|---|---|---|---|---|---|---|
| ASO2 | 803.5 | 95.0 | 50 | -90 | -10 | -120 | -25 |
| ASO4 (IS) | 896.0 | 95.0 | 50 | -110 | -10 | -115 | -15 |

### FIGURES:

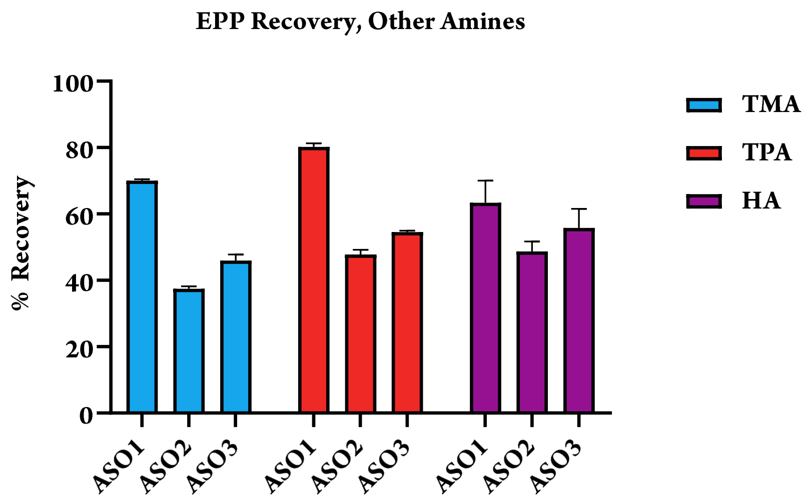

**Supporting Figure S.1. Assessment of recoveries for ASO1, ASO2 & ASO3 using other small volatile amines such as Trimethylamine (TMA), Tripropylamine (TPA) & Hexylamine (HA) instead of ammonia in extraction solution.**

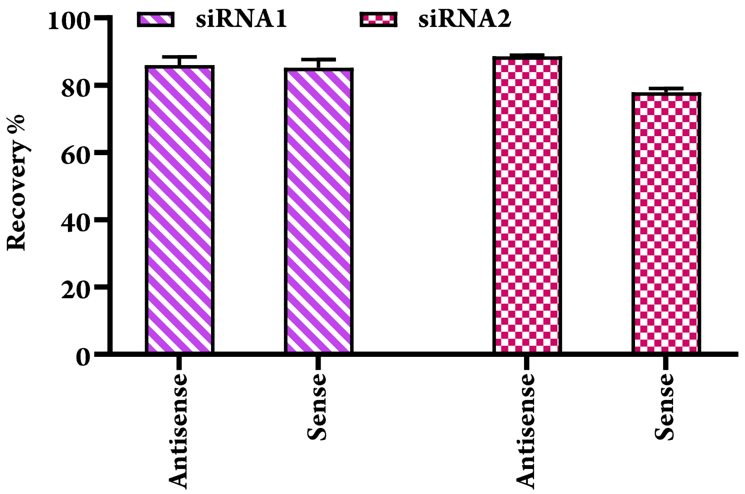

**Supporting Figure S.2. EPP extraction recovery for both antisense and sense strands of GalNAc-conjugated siRNA1 and Lipid-conjugated siRNA2**

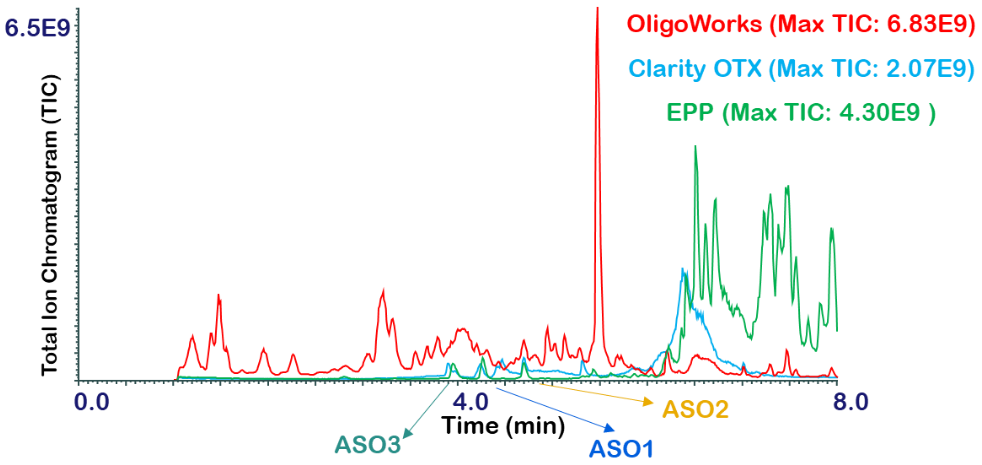

**Supporting Figure S.3. Comparing total ion chromatograms across the three extraction methods to illustrate the LCMS background signal for each method.**

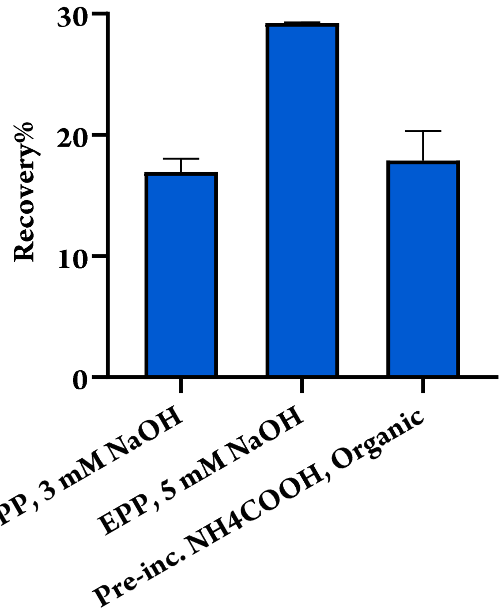

**Supporting Figure S.4. Assessing the effect of using strong base (sodium hydroxide) in extraction solution and comparing ionic reagents (ammonium formate and sodium chloride) with ammonia via pre-incubation in the sample; on the recovery of ASO1.**

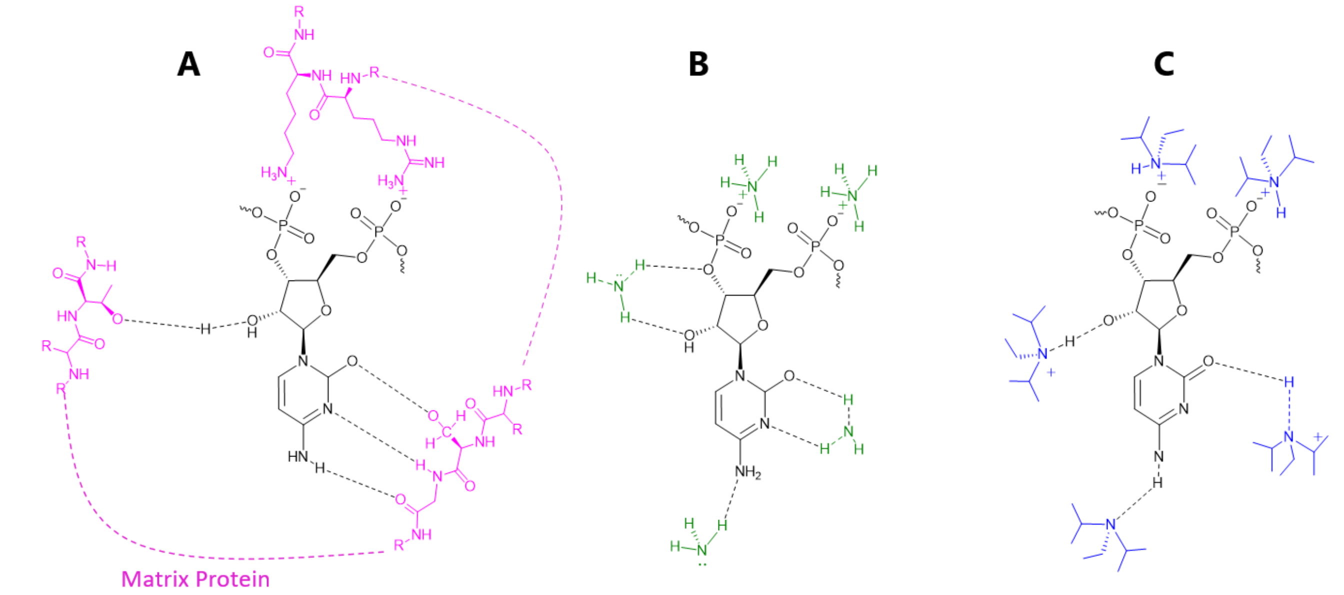

**Supporting Figure S.5. (A) Illustration of possible nucleotide interactions with amino acid residues of biological matrix proteins via hydrogen bond and ionic interactions. (B) Illustration of ammonia forming ionic interactions and hydrogen bonds with nucleotides. (C) ionic and hydrogen bond interactions of DIPEA with the nucleotide.**
